# Feasibility of Ultra-Short Term Analysis of Heart Rate and Systolic Arterial Pressure Variability at Rest and During Stress via Time-domain and Entropy-based Measures

**DOI:** 10.1101/2022.10.29.514356

**Authors:** Gabriele Volpes, Chiara Barà, Alessandro Busacca, Salvatore Stivala, Michal Javorka, Luca Faes, Riccardo Pernice

## Abstract

Heart Rate Variability (HRV) and Blood Pressure Variability (BPV) are widely employed tools for characterizing the complex behavior of cardiovascular dynamics. Usually, HRV and BPV analyses are carried out through short-term (ST) measurements, which exploit ∼5 minute-long recordings. Recent research efforts are focused on reducing the time series length, assessing whether and to what extent Ultra-Short Term (UST) analysis is capable of extracting information about cardiovascular variability from very short recordings. In this work, we compare ST and UST measures computed on electrocardiographic R-R intervals and systolic arterial pressure time series obtained at rest and during both postural and mental stress. Standard time-domain indices are computed, together with entropy-based measures able to assess regularity and complexity of cardiovascular dynamics, on time series lasting up to 60 samples, employing either a faster linear parametric estimator or a more reliable but time-consuming model-free method based on nearest neighbor estimates. Our results evidence that shorter time series up to 120 samples still exhibit an acceptable agreement with the ST reference, and can be exploited to discriminate between stress and rest as well. Moreover, although neglecting nonlinearities inherent to short-term cardiovascular dynamics, the faster linear estimator is still capable of detecting differences among the conditions, thus resulting suitable to be implemented on wearable devices.

## 1. Introduction

In recent years, an increased interest is being devoted to the study of the complex regulatory mechanisms of the human organism, in order to improve the level of knowledge of biological functions and the early detection of pathologies [1–3]. In particular, a large amount of information can be achieved studying the cardiovascular system, both alone and through its interactions with other physiological systems, by analyzing several parameters, such as blood pressure, level of oxygenated hemoglobin, heart rate and heart rate variability (HRV) [4–6]. The latter represents the beat-to-beat variation of the duration of the cardiac cycle, and allows to obtain information not only about the cardiovascular function, but also about the balance between the activities of the sympathetic nervous system (SNS) and the parasympathetic nervous system (PNS), thus allowing a thorough understanding of the complex neuro-autonomic regulation. This is important because the usually antagonistic actions of the SNS and the PNS vary in response to psycho-physical stress situations that can occur in different contexts during daily life of both healthy and pathological individuals [7,8]. Similarly, the study of other cardiovascular parameters, such as systolic (SAP) and diastolic (DAP) arterial pressure variability, enables to investigate the multiple feedback (e.g., baroreflex control) and autoregulatory mechanisms (e.g., vascular myogenic autoregulation) that come into play in the complex cardiovascular regulation [9–14].

HRV is usually studied through the monitoring of electrocardiographic (ECG) recordings, extracting the time series of R– R intervals (i.e., the time periods between successive heartbeats) [15]. In clinical settings, the use of 24-hour recordings, also referred as long term (LT) analysis, is considered the gold standard for the investigation of cardiovascular control mechanisms, since such timeframe allows a better description of the physiological processes, taking into account slower temporal fluctuations (e.g., the circadian rhythms), and the response of the organism to a wider range of external stimuli as well [11,15]. On the other hand, short-term (ST) measurements are typically based on 5 minutes recordings and have been widely employed especially for assessing the balance between SNS and PNS activities, given that fluctuations mediated by autonomic nervous system (ANS), reflecting respiratory, baroreflex and vascular tone regulatory mechanisms overlap to generate short-term dynamics [9–11,15]. Similarly, also blood pressure variability (BPV) can be studied through short-term and long-term analyses; its variations have been shown to be the result of complex interactions between extrinsic environmental factors and intrinsic cardiovascular regulatory mechanisms and have been associated to the risk of cardiovascular events and mortality [13,14,16].

Short-term HRV is commonly investigated through different time-, frequency- and information-theoretic domain indexes computed starting from ECG R-R interval time series. Specifically, time-domain indexes are used to quantify both average heart rhythm and the extent of beat-to-beat variability [15,17], while frequency-domain indexes extract information specific to various time scales of oscillations; furthermore, more recently developed entropy-based measures permit to assess the regularity and complexity of cardiovascular dynamics [18–24]. ST HRV analysis has been proven useful also outside clinical settings, e.g. to monitor health and wellbeing at home and in everyday life scenarios using wearable smartwatches and smartbands [25–28].

With the widespread adoption of wearable biomedical devices especially in domestic settings (e.g. smart-healthcare), the research is now focusing on whether and to what extent shorter recordings can be exploited for cardiovascular variability analysis, given their lower computational and memory resources, in order to quickly extract useful physiological indexes [25,26,29,30]. Several works have recently investigated on the so-called ultra-short term (UST) HRV analysis which exploits recordings shorter than 5 minutes, comparing the results with those obtained using the ST gold standard [31–36]. However, the choice of the time series length strongly influences the physiological indices derived from RR and blood pressure time series, in a way such that employing shorter recordings reduces the ability to resolve slower oscillations within the analyzed cardiovascular dynamics [17,37]. Generally, at least 2-minute recordings are recommended to observe Low-Frequency (LF, range: 0.04-0.15 Hz) dynamics (related especially to SNS, but also to PNS activity) and at least 1-minute to observe High-Frequency (HF, range: 0.15-0.4 Hz) dynamics (mainly related to parasympathetic activity fluctuations associated with respiration) [15,17]. Longer recordings, i.e. 24-hour period LT analysis, further allow to detect lower frequency components, such as the very-low-frequency (VLF, 0.0033-0.04 Hz) and the ultra-low-frequency (ULF, <0.003 Hz). Therefore, the use of recordings shorter than 5 minutes may bring a loss of information related to slower dynamics if compared to ST analysis. The reliability of the cardiovascular parameters computed from UST recordings also depends on the acquisition protocol and on the dynamics of the response mechanisms to the task or stimulus [37].

The present work aims at evaluating the extent to which the loss of physiological information due to UST reduced time series length can be a good tradeoff for extracting physiological indices with lower real-time processing and storage costs, suitably for wearable devices [28,31,36,38–40]. Moreover, while several works have focused on ultra-short term HRV [31,36,40,41], to the best of our knowledge there are no previous studies performing an UST blood pressure variability analysis. Herein, a comparison between UST and ST indices extracted in the time and information domains is performed on a dataset composed of systolic blood pressure (SAP) and inter-beat interval (RR) time series acquired on a population of healthy subjects in rest and when undergoing to orthostatic and mental stress. The analysis has been carried out reducing the time series length from 300 (short-term) to 60 samples, in steps of 60, to assess the loss of information at decreasing window length and to verify whether the shortest length is still able to discriminate the transition from rest to stress.

## 2. Materials and Methods

### 2.1 Experimental protool

Analyses were carried out on an historical dataset employed for assessing the effects of orthostatic and mental stress on cardiovascular dynamics. Data have been acquired on 61 healthy young volunteers (24 males, 37 females) aged 17.5 years ±2.4 years, normotensive and having a normal body mass index (BMI=19÷25 *kg m*^−2^) [20,42]. All participants signed a written informed consent before taking part in the measurement protocol, also requiring a parental or legal guardian permission to participate in the study when the subject was a minor (i.e., less than 18 years of age). All procedures were approved by the Ethical Committee of the Jessenius Faculty of Medicine, Comenius University, Martin, Slovakia. Subjects were asked not to take substances influencing the autonomic nervous system and cardiovascular system activities [20,42].

Physiological signals recorded on the volunteers consisted of (i) electrocardiographic (ECG) signal acquired through a horizontal bipolar thoracic lead (CardioFax ECG-9620, NihonKohden, Japan), (ii) continuous arterial blood pressure (BP) recorded on the finger through the volume-clamp method (Finometer PRO, FMS, Netherlands). All signals were acquired synchronously with a sampling frequency of 1 kHz.

Subjects were positioned on a motorized tilt table and a restraining strap at the thigh level was put to ensure safety and stability of the subject during the movement of the tilt table. Signals were acquired during a measurement protocol consisting of the following four phases (schematically represented in Fig.1 (a)):

- A resting condition (R1) with the subject laying in the supine position for 15 minutes, in order to stabilize the physiological signals on a baseline level;
- A head-up tilt (T) test aimed at evoking mild orthostatic stress by inclining the motorized table of 45 degrees for 8 minutes;
- Another resting condition (R2) with the subject laying in the supine position for 10 minutes, in order to restore the physiological parameters to their baseline values;
- A 6-min long mental arithmetic (M) task aimed to evoke cognitive load (i.e., mental stress), during which subjects were asked to mentally carry out the sum of three digits in the least possible time, indicating whether the result was an even or odd number.

**Figure 1.**
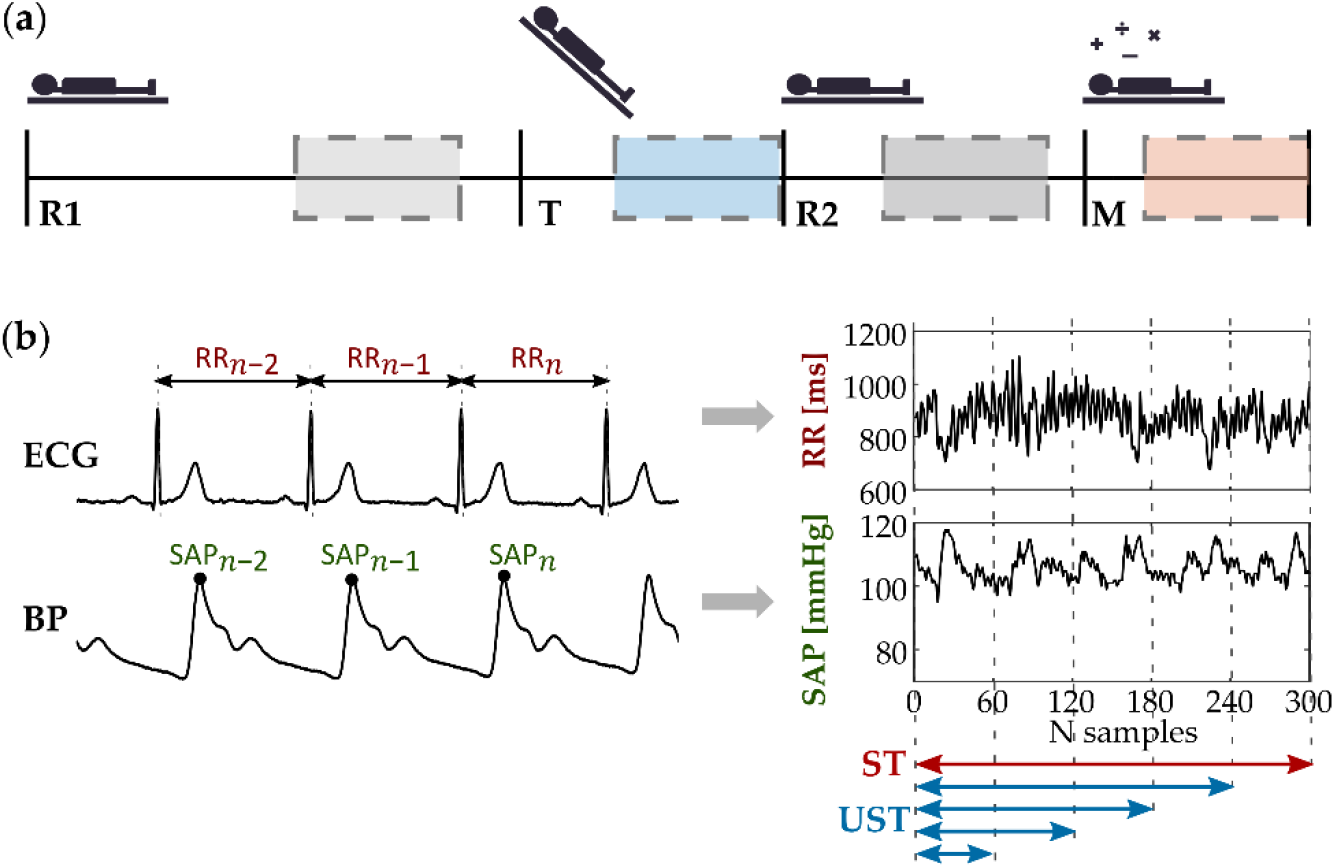
**(a)** Schematic illustration of the experimental protocol, including baseline resting (R1), orthostatic stress (T), second resting (R2) and mental stress (M). **(b)** Representative RR and SAP time series, extracted respectively from ECG and BP recordings, which have been investigated through univariate analysis performed after short-term (ST, 300 points, in red) and ultra-short term (UST, 240 to 60 points, in blue) time window segmentation.

During the whole measurement protocol, the subjects were asked to avoid any movement or speaking, to decrease artifacts occurrence and minimize the non-stationarities during recording of the signals.

### 2.2 Time series extraction

Starting from the ECG and BP signals, acquired for each subject and condition we have extracted time series for carrying out the analyses described in the following subsections. Specifically, the RR interval time series was obtained measuring the temporal distance between consecutive ECG QRS apexes, while the systolic arterial pressure (SAP) series was obtained as the sequence of the maximum values of the blood pressure signal measured within each RR interval.

Initially, time series of 300 heartbeats were extracted according to the standard of short-term analysis. The analyzed windows started 8 min after the beginning of the R1 phase, 3 min after the beginning of the T phase, 3 min after the beginning of the R2 phase and 2 min after the beginning of the M phase (see schematization in Fig. 1(a)). This choice has been done in order to favor the stationarity of the time series, neglecting the transition effects due to the physiological changes elicited by the different phases during the measurement protocol [20,42]. Before performing the analyses, a visual inspection of the series was carried out to check for their stationarity.

Afterwards, in order to perform ultra-short term (UST) analysis of cardiovascular parameters, shortened time series were obtained by reducing the series length each time of 60 samples up to a minimum of 60 heartbeats. The resulting UST time series were composed of 240, 180, 120 and 60 samples, selected starting from the beginning of the reference ST series, as schematized in Fig. 1(b).

### 2.3 Time domain analysis

Time domain analysis was performed on ST and UST RR and SAP time series computing the average (MEAN) and the standard deviation (STD) of the series values. With regard to RR time series, the standard deviation coincides with the widely used standard deviation of the interbeat intervals between normal sinus beats (SDNN) [15], given that no ectopic beats were detected. Moreover, for the RR time series, the root mean square of successive differences (RMSSD) was computed as follows to extract information about the beat-to-beat changes in heart rate mediated mostly by PNS [15,23,43]:

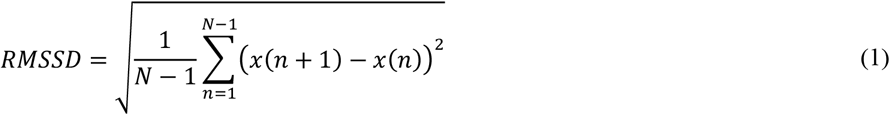

being *x*(*n*) the *n*-th RR samples and *N* the time series length.

### 2.4 Information domain analysis

For both RR and SAP time series, information-theoretic analysis was performed to quantify the information carried by the physiological time series, as well as their complexity. The latter is typically quantified as the unpredictability of the present sample given its past samples, and thus has been associated to the regularity of the time series [21,44]. For this reason, in this work the static entropy (SE), dynamic entropy (DE) and conditional entropy (CE) measures were computed on either short-term or ultra-short-term series using both a parametric and a model-free estimation.

Starting from a stationary stochastic process *X*, let us denote as *x* = {*x*_1,_ *x*_2,_ …, *x*_*N*_} the time series of length *N*, taken as a realization of the process *X*, as *X*_*n*_ the variable obtained by sampling the process *X* at the present time *n* and 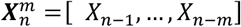 the variable describing the collection of the past *m* states. Using such notation, the static entropy quantifies the ‘static’ information contained in the current state of the process *X*, without considering its temporal dynamics, and can be defined as [45]:

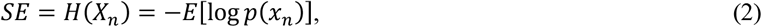

where *E*[·] is the expectation operator and *p*(·) the probability density, while *H*(·) denotes the entropy. The dynamic entropy (DE) instead represents the “joint” entropy of the present and past variables composing the process; therefore it provides the amount of information brought by the current sample of the series and by its past samples as well, thus giving ‘dynamic’ information on the entire process, and it can be defined as [46]:

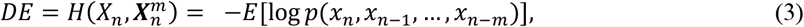

being *H*(·,·) the joint entropy of two random variables. Then, the conditional entropy (CE) quantifies the average uncertainty that remains about the present state of the process when its past states are known (i.e., the new information contained in the current sample that cannot be inferred from the past history), and is defined as [45]:

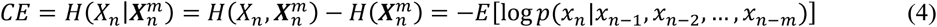

where *H*(· | ·) denotes conditional entropy operator.

In this work, the computation of SE, DE and CE indices was carried out through two different estimation approaches, in order to identify which method allows the best trade-off between computational costs and ability to discriminate physiological changes (i.e. rest versus stress).

The first estimation method (hereinafter referred as *lin*) consists of a linear parametric approach based on the assumption that the observed process *X* is a stationary Gaussian process [45], which is a reasonable assumption given that many physiological data tend to a Gaussian distribution. Under this assumption, the above-mentioned entropies measures can be computed, after describing the dynamics of the process *X* with a linear regression model, from the covariance matrices of the variables sampling the process. In particular, the present and past variables of the process are related with the autoregressive (AR) model 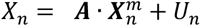, where ***A*** is a vector of *m* regression coefficients and *U* is a white noise process modeling the prediction error. AR model identification has been performed via ordinary least squares method [47] to obtain estimations of regression parameters and prediction error variance, thus estimating the variance and the covariance matrices of the process. Then, denoting as 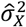 the variance of the process, as 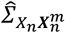 the covariance matrix of the present and past states of *X*, and as 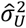 the prediction error variance, the above defined entropy measures can be computed as [46]:

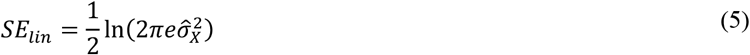

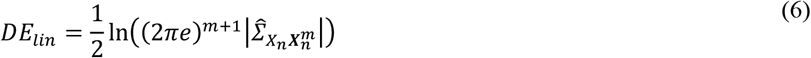

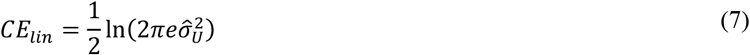

The second estimation method (hereinafter referred as *knn*) is a model-free approach based on nearest neighbor metrics, which exploits the intuitive notion that the local probability density around a given data point is inversely related to the distance between the point and its neighbors. Using this approach, estimates of SE, DE and CE of the process *X* can be respectively computed through the following expressions [45,46]:

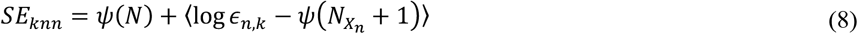

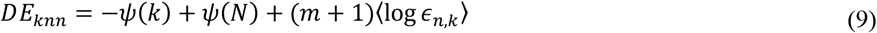

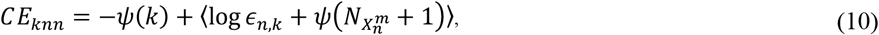

where *ψ*(·) is the digamma function, *k* is the number of neighbors chosen for the analysis, *ϵ*_*n,k*_ represents twice the distance between the *n*-th realization of 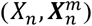and its *k*-th nearest neighbor, 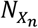 and 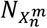 respectively represent the number of points with a distance from *x* and 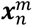 smaller than 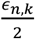 and ⟨·⟩ is the average operator; the average is taken over all the *N* − *m* realizations of the patterns 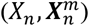that can be extracted from a series of length *N*. Here, estimation of the SE and CE in (8) and (10) has been performed exploiting the distance projection method for bias compensation described in [45,46].

The ECG and SBP signals were pre-processed and analyzed using MATLAB R2021b (The MathWorks, Inc.). All the RR and SAP time series were normalized to zero mean before computing the three entropy measures. For computing DE and CE the time series were also normalized to unit variance; this was not carried out for static entropy, whose calculation requires the knowledge of the information of the variance of the time series [46]. Information-domain indices were computed using the online available Matlab ITS Toolbox (see Data availability statement section).

The number of neighbors chosen for the model-free estimator was *k*=10, while the number of past components considered for the time series past histories was set equal to 2. Similarly, the order of the autoregressive model defined using standard least-squares regression for the *lin* approach was set to *m*=2.

In order to quantify the computational resources required to compute these indices, the time spent for calculating linear and model-free estimations of DE, CE and SE was measured for both RR and SAP time series in the four considered phases at varying time series length using the built-in MATLAB function.

### 2.5. Statistical analysis

The statistical analyses have been carried out on distributions of time-domain and entropy measures obtained in the four phases (R1, T, R2, M) on both RR and SAP time series. Given that the normality of distributions for the analyzed indexes was verified according to the Kolmogorov-Smirnov test, the parametric Student’s t-test was used to perform the pairwise comparisons, with a significance threshold set to *p* < 0.05. Specifically, the statistical tests were carried out to compare (i) orthostatic and mental stress conditions with resting states (i.e., T vs R1 and M vs R2) and (ii) ultra-short-term and short term distributions (i.e., UST vs. ST).

However, the mere use of statistical tests has been often considered not sufficient for assessing feasibility of using HRV indices evaluated through different techniques in studies where it has been supported by other approaches, e.g. correlation analysis, Bland–Altman plots or effect size [23,36,48,49]). In our work, correlation analysis was carried out through the computation of the squared Pearson correlation coefficient *r*^2^ to obtain a measure of the agreement between each UST and the ST reference distribution. According to Shafer *et al*. [40,50] who selected a conservative criterion for the Pearson correlation coefficient (*r* ≥ 0.90), herein we set a threshold for the squared coefficient equal to *r*^2^ = 0.81 to establish the presence of a strong agreement between indexes derived from UST and ST analysis. Moreover, for all the time and information domain indices and time series length, we assessed the difference between the distributions during stress and rest (i.e., T vs R1 and M vs R2) by computing the “effect size “through the Cohen’s *d* measure [51]:

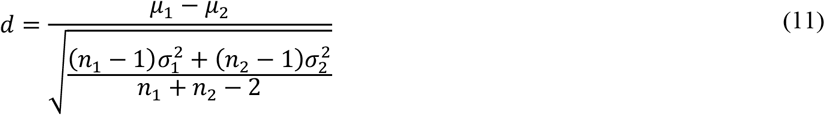

where *μ, σ* and *n* are respectively the mean, the standard deviation and the number of samples of the two distributions under comparison (i.e., the number of subjects). Generally, the effect size is deemed as small, medium and large, if the absolute value of *d* is lower than 0.2, between 0.2 and 0.5 or higher than 0.8, respectively.

## 3. Results

### 3.1. Time domain analysis

Figure 2 shows the comparison of ST (N=300) and UST (N<300) analyses performed for the time domain indexes (MEAN, SDNN, RMSSD) computed over the RR time series across the 61 subjects in the four considered phases. For each panel, the top row subplot shows the boxplot distributions of the indices, the central one the Cohen’s *d* measure (in absolute value) and the bottom one the Pearson squared correlation coefficient. Results show a statistically significant decrease of all the three indexes during T vs R1 and during M vs R2 phases for the ST and for all the UST time window lengths taken into account. With regard to MEAN, statistically significant differences between UST and ST distributions are reported only for R2 already from *N*=240 and for shorter time series (Fig. 2(a.2), top subplot). As regards SDNN, statistically significant differences have been detected comparing all the UST distributions to ST reference during head-up tilt (Fig. 2(b.1), top subplot). On the contrary, statistically significant differences have been reported only in the shortest UST distribution (N=60) for both R2 and M (Fig. 2(b.2, top subplot)). No statistically significant differences have been reported for RMSSD. Cohen’s *d* measures (central subplots in Fig. 2) computed between stress and rest reported a medium-to-high effect size (|*d*| **≥** 0.7) for all the three indices, but lower for RMSSD during mental stress (|*d*|≈0.5). Moreover, in all the cases the Cohen’s *d* showed higher values with regard to postural stress discrimination rather than mental stress. Furthermore, *d* remains almost constant at decreasing time series length, except for SDNN assessed during mental stress (Fig. 2(b.2)), in which it decreases with *N*. The squared Pearson correlation coefficient (bottom panels in Fig. 2) computed between ST and UST distributions is very high and almost always above the threshold (*r*^2^ > 0.81) for MEAN and RMSSD indices for all the time window lengths, except for RMSSD during T for *N*=60. A considerable decrease of correlation is reported for SDNN index, going below threshold for T when N=120 (Fig. 2(b.1, bottom panel)), and for both R2 and M when N=60 (Fig. 2(b.2, bottom panel)).

**Figure 2.**
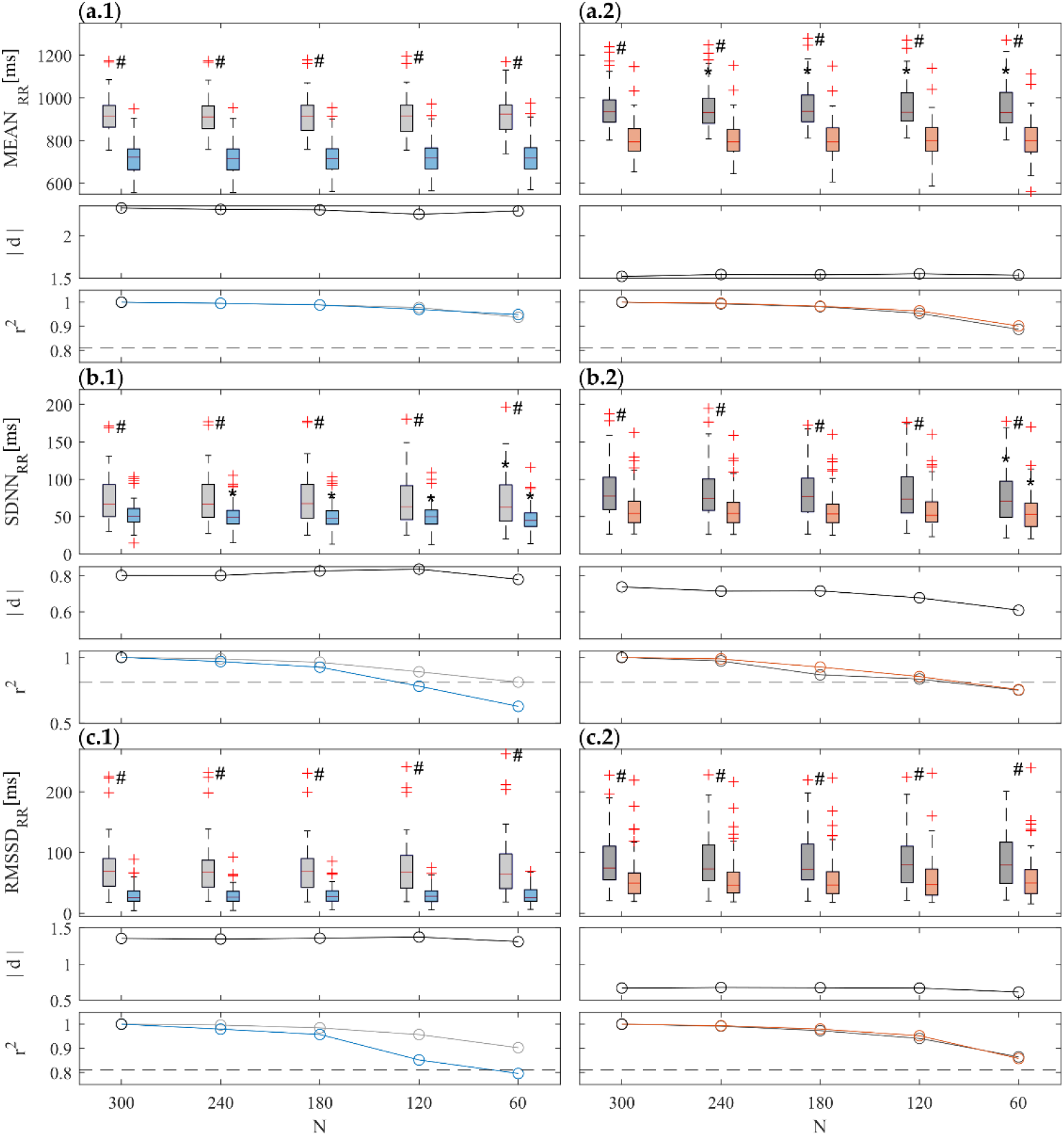
Boxplot distributions (top subplots) of time domain indexes, i.e. (**a**) MEAN, (**b**) SDNN and (**c**) RMSSD calculated from RR time series during R1 (light gray) and T (light blue), (.**1** panels), and during R2 (dark gray) and M (orange) (.**2** panels) phases. Statistical tests: #, p<0.05, T vs R1 or M vs R2; *, p<0.05, ST vs UST. Central subplots: Cohen’s *d* (in absolute value) evaluated between each stress condition and the previous rest phase (i.e., R1-T and R2-M) for all the considered time series lengths. Bottom subplots: squared Pearson correlation coefficients computed between a given UST distribution and the ST reference, with a threshold of r^2^ = 0.81 (dotted gray line).

Figure 3 shows the comparison of ST (N=300) and UST (N<300) analyses with regard to the time domain indexes (MEAN, STD) of SAP time series across the 61 subjects in the four considered phases. With regard to ST, MEAN decreases significantly during T if compared to R1 (Fig. 3(a.1), top subplot), while increases significantly during M if compared to R2 (Fig. 3(a.2), top subplot); opposite trends are reported with regard to STD (Fig. 3(b.1-b.2), top subplots). For both indexes, results show statistically significant differences for T vs R1 and for M vs R2 phases for all the UST time window lengths taken into account, except for T vs R1 with regard to the shortest series length (N=60) only for STD (Fig. 3(b.1, top subplot)). With regard to MEAN, statistically significant differences have been reported comparing UST vs ST distributions during head-up tilt condition for N=240 and shorter (Fig. 3(a.1), top subplot)); on the other hand, no statistically significant differences have been reported with regard to R1 and to both M and R2 conditions (Fig. 3(a.2, top subplot)). As regards STD, statistically significant differences have been reported comparing UST vs ST distributions for N **≤** 120 and N **≤** 180, respectively for R1 and T (Fig. 3(b.1, top subplot)), for N **≤** 240 for R2 and for just N=60 for M (Fig. 3(b.2, top subplot)). Cohen’s *d* values evidence a high effect size (|*d*| > 0.8), except for STD index during R1-T transition (Fig. 3(b.1, central subplot)) in which there is a medium-low effect size (|*d*| ≈ 0.5). An overall decrease in effect size is observed with the sample size *N*, especially for STD. The correlation analysis between UST and ST distributions reported a high squared correlation coefficient (r^2^ > 0.81) with regard to MEAN distributions (Fig. 3(a.1) and (a.2), bottom subplots). On the other hand, as regards to STD, the correlation coefficient strongly decreases when shortening *N*, going below the threshold for N=120 for R1, R2 and M and for N=60 for T (Fig. 3(b.1) and (b.2), bottom subplots)).

**Figure 3.**
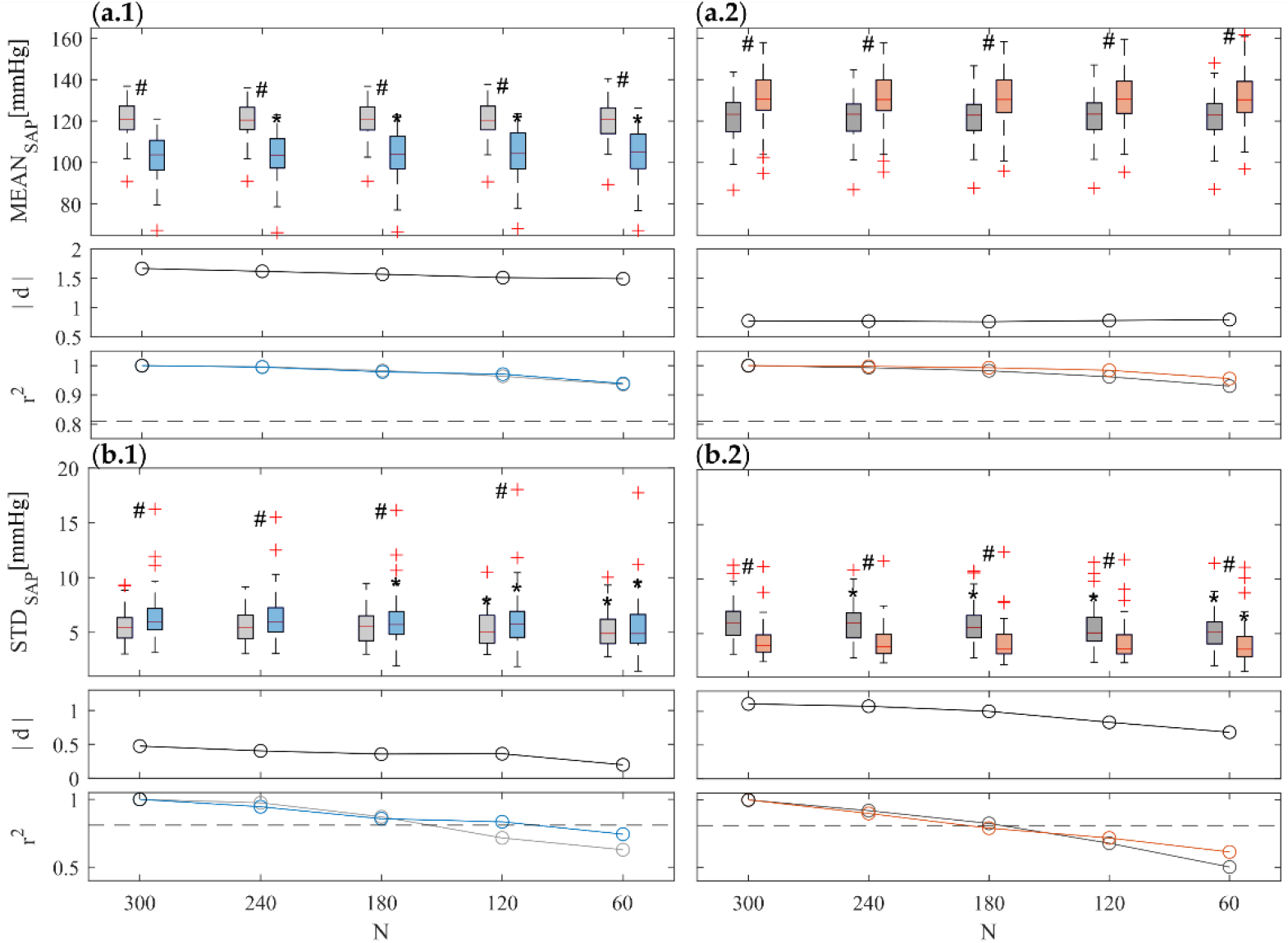
Boxplot distributions (top subplots) of time domain indexes, i.e. (**a**) MEAN and (**b**) STD calculated from SAP time series during R1 (light gray) and T (light blue), (.**1** panels), and during R2 (dark gray) and M (orange) (.**2** panels) phases. Statistical tests: #, p<0.05, R1 vs T and R2 vs M; *, p<0.05, ST vs UST. Statistical tests: #, p<0.05, T vs R1 or M vs R2; *, p<0.05, ST vs UST. Central subplots: Cohen’s *d* (in absolute value) evaluated between each stress condition and the previous rest phase (i.e., R1-T and R2-M) for all the considered time series lengths. Bottom subplots: squared Pearson correlation coefficients computed between a given UST distribution and the ST reference, with a threshold of r^2^ = 0.81 (dotted gray line).

### 3.2 Information domain analyses

Figures 4 and 5 depict the results of the information domain analysis carried out computing SE, DE and CE indices through both *lin* and *knn* estimators respectively for RR and SAP time series, across the 61 subjects for each of the four physiological conditions (R1, T, R2 and M). For each panel, the top row subplot shows the boxplot distributions of the indices, the central one the Cohen’s *d* measure (in absolute value) and the bottom one the Pearson squared correlation coefficient.

**Figure 4.**
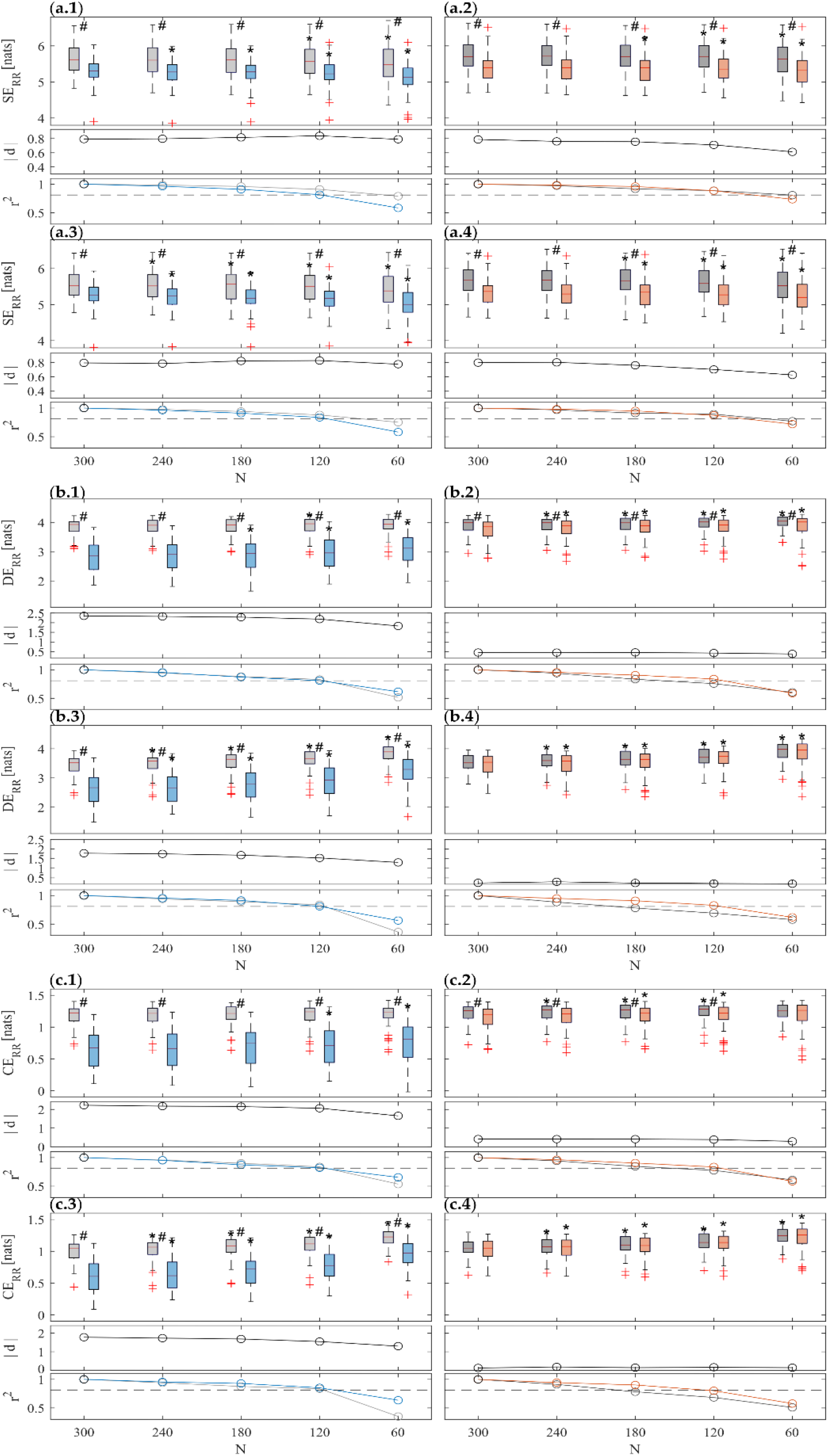
Results of information domain analysis on RR time series. Boxplot distributions (top subplots) of ST and UST indices of (**a**) SE, (**b**) DE and (**c**) CE calculated using both *lin* (.**1** and .**2**) and *knn* (.**3** and .**4)** estimators during R1 (light gray) and T (light blue) (.**1** and .**3**), and during R2 (dark gray) and T (orange) (.**2** and .**4**) phases. Statistical tests: #, p<0.05, T vs R1 or M vs R2; *, p<0.05 ST vs UST. Central subplots: Cohen’s *d* (in absolute value) evaluated between each stress condition and the previous rest phase (i.e., R1-T and R2-M) for all the considered time series lengths. Bottom subplots: squared Pearson correlation coefficients computed between a given UST distribution and the ST reference, with a threshold of r^2^ = 0.81 (dotted gray line).

**Figure 5.**
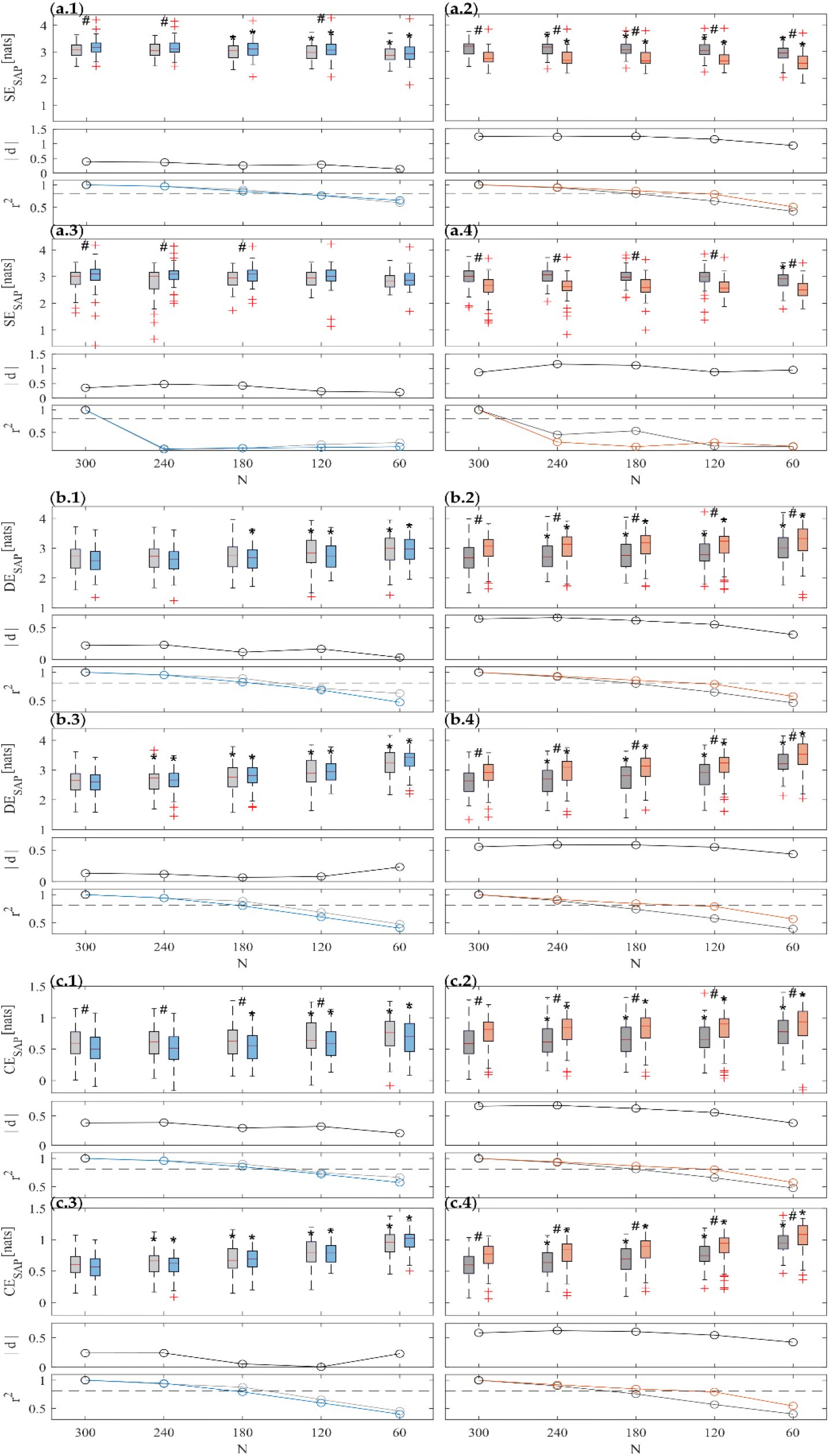
Results of information domain analysis on SAP time series. Boxplot distributions (top subplots) of ST and UST indices of (**a**) SE, (**b**) DE and (**c**) CE calculated using both *lin* (.**1** and .**2**) and *knn* (.**3** and .**4)** estimators during R1 (light gray) and T (light blue) (.**1** and .**3**), and during R2 (dark gray) and T (orange) (.**2** and .**4**) phases. Statistical tests: #, p<0.05, T vs R1 or M vs R2; *, p<0.05 ST vs UST. Central subplots: Cohen’s *d* (in absolute value) evaluated between each stress condition and the previous rest phase (i.e., R1-T and R2-M) for all the considered time series lengths. Bottom subplots: squared Pearson correlation coefficients computed between a given UST distribution and the ST reference, with a threshold of r^2^ = 0.81 (dotted gray line).

The results of the analyses carried out on RR time series are shown in Fig. 4. For both orthostatic and mental arithmetic conditions, the shift from rest to stress highlights significant decrease of SE, DE and CE measures computed with both *lin* and *knn* estimators. This variation is most appreciable under the postural stress, in fact statistically significant variations are always reported comparing T vs R1, for both ST and all the UST distributions and for both estimators. On the other hand, using the *lin* estimator statistically significant variations are reported comparing M vs R2, for both ST and almost all the UST distributions for all measures (except w.r.t to CE for N=60), but only for SE (Fig. 4(a.4)) using the *knn* approach. SE appears to decrease while CE and DE to increase when decreasing the time series length N. This result is more evident using the *knn* estimator, in fact statistical analysis carried out between UST and ST distributions highlighted significant differences starting from time series of length N=240 for almost any measure computed through the model-free approach. For DE and CE computed through *lin* estimator, the statistically significant differences between UST and ST at T occur only for N **≤** 180 and N**≤** 120, respectively. The Cohen’s *d* values obtained for *lin* estimator are higher than for the *knn* one (see central subplots in each panel in Fig. 4). High effect sizes are reported for CE and DE during T, but medium values instead during M; with regard to the SE index, a medium-high effect size is assessed for both phase transitions and for both estimators. In any case, *d* appears almost constant at decreasing N (up to N=120), while a more marked decrease is observed when going to N=60. Finally, the squared Pearson correlation coefficient (see bottom subplots in each panel in Fig. 4) decreases shortening the time series length N, still largely showing a high degree of correlation (above the threshold) up to N=120.

As regards entropy measures computed on the SAP time series reported in Figure 5, both estimators (i.e., *lin* and *knn*) and analysis approaches (i.e., ST and UST) show an increase of SE (Fig. 5(a.1, a.3), top subplots) as well as a decrease of DE (Figs. 5(b.1, b.3), top subplots) and CE (Figs. 5(c.1, c.3), top subplots) from R1 to T, and conversely a decrease of SE (Fig. 5(a.2, a.4), top subplots) and an increase of DE (Figs. 5(b.2, b.4), top subplots) and CE (Figs. 5(c.2, c.4), top subplots) from R2 to M. Nevertheless, while differences between M and R2 distributions are always statistically significant, the comparison between T and R1 evidenced statistical significance only for *lin* estimation of CE using time series not shorter than 120 samples and for SE obtained with both estimators and N **≤** 240.

As regards the comparison between ST and UST analyses, for both estimators and phase shifts, CE and DE values increase as the time series length decreases, whereas the SE decreases more slowly. Statistical analysis highlighted significant differences already from the first window length (N=240) for almost all the information indices obtained with both estimators during R2 and M, while overall this is true for T vs R1 in any cases only for shorter time series (N<180). The only exception is the *knn* estimation of SE, for which no significant differences are found between ST and UST analysis. The effect size assessed through Cohen’s *d* (Fig. 5, central subplots in each panel) is always medium-low, except for SE index for M vs R2, and overall decreases in absolute value when shortening the series length. The correlation analysis between UST and ST distributions (Fig. 5, bottom subplots in each panel) shows that the squared Pearson’s correlation coefficient decreases when reducing time series length, still reporting values higher than the threshold (*r*^2^ = 0.81) for almost all indices for N>=180 (often even for N=120), except for the SE estimated with the *knn* approach for which *r*^2^ severely drops already for N=240 (Figs. 5(a.3, a.4), bottom subplots).

Finally, we report the results relevant to the computational times required for the calculation of the entropy-based measures, performed using both estimators. In order to compare ST and UST analysis times, we have selected N=120 samples as UST time series length, since the previous results highlighted that this is the minimum length which overall guarantees a very good agreement between ST and UST distributions. The average computation time of all the entropy measures on 488 iterations (2 time series in 4 different physiological conditions for 61 subjects) was 0.24 ms and 5.87 ms respectively for *lin* and *knn* on ST 300-samples time series, and 0.17 ms and 1.90 ms on UST 120-samples time series. Such computational times were obtained on a computer equipped with an Intel Core i7-11700K CPU (3.60 GHz), 64 GB RAM, 512 GB SSD, Windows 11, MATLAB R2021b. The computational times are similar for RR and SAP series and do not vary as well with the protocol phase. Moreover, while computational times remain almost constant as time series length decreases with regard to *lin*, they strongly decrease at shortening N with regard to model-free estimation.

## 4. Discussion

The aim of this work was to evaluate to what extent UST HRV and BPV measurements of less than 5-min duration can be used as a substitute of the widely employed and validated short-term recordings. Employing standard time-domain and recently introduced entropy-based measures, we aim to assess whether and which UST metrics can replace ST indices to discriminate postural and mental stress states, while exploiting shorter recordings that are less prohibitive in terms of time cost and computational power. In this sense, we also compared two different approaches for estimating entropy-based measures, i.e. a more reliable but also more computationally intense non-linear model-free method and a faster but less general model-based approach. The rest of the Discussion is organized as follows. Subsections 4.1 and 4.2 focus on physiological interpretation of results obtained through ST time-domain and information-theoretic domain metrics, respectively. Subsection 4.3 instead discusses UST results in terms of agreement with standardized ST measures.

### 4.1 Time domain analyses on ST series

Time-domain HRV results (Fig. 2) are in agreement with the widely recognized findings in the literature which evidence an increased heart rate, a decrease of variability (SDNN) and of RMSSD during stress conditions, in particular after head-up tilt [23,52,53]. In both physiological stress conditions, but less markedly during mental stress, these trends are related to an enhanced sympathetic and reduced parasympathetic activity, i.e. resulting from a SNS activation and a PNS withdrawal, resulting in a shift in sympathovagal balance [54–56]. In particular, the reduced parasympathetic contribution is evidenced by the decreased RMSSD which has been usually related to PNS activity [15,17]. Nevertheless, physiological mechanisms involved during orthostatic and arithmetical stress states are different, as demonstrated by different SAP MEAN and STD trends in these two stress conditions (Fig. 3). This is in agreement with previous studies highlighting the presence of a closed-loop regulatory mechanism between RR and SAP related to mechanical and baroreceptive actions of the vascular system that, together with ANS activity, is responsible for changes in cardiovascular dynamics [10,57,58]. The decrease of SAP average together with the increase of its variability during postural stress (see Fig. 3(a.1,b.1)) have been related to the decreased venous return [59–61]. The resulting cardiac filling associated with SAP decrease leads to an activation of the baroreflex response during postural stress and thus to an enhanced involvement of baroceptive mechanism and vasoconstrictor activity in response to the physiological perturbation, which in turn produces an increased heart rate [59–62]. The opposite trends reported for mental stress (see Fig. 3(a.2,b.2)) can be put into relation to cortical mechanisms related to mental stress eliciting vasomotor reactions reflected in SAP changes [53,63,64].

### 4.2 Information domain analyses on ST series

The shift of the autonomic balance to the sympathetic branch caused by orthostatic and cognitive challenges produces a simplification of the cardiac dynamics, with a reduced information contained in the RR time series (see Fig. 4(a.1-a.4)), which has been put into relation to the dominant oscillations centered around the frequency of Mayer waves [20,23]. The elicited stress conditions lead also to a decrease in complexity, and thus lower CE and DE values using both linear and non-linear estimators (see Fig. 4(b.1-c.4)); physiologically, this indicates a regularizing effect on the cardiac dynamics produced by sympathetic activation and vagal withdrawal already demonstrated in several previous works also on the same dataset [21–23,44,65]

The entropy-based SAP analysis revealed opposite trends for T and M compared to the preceding resting condition (see Fig. 5), confirming once again the different response to postural and mental stress. Mental challenge produced a SE decrease and increased complexity, while opposite trends have been observed for postural stress, even if they are significant only using *lin* estimator. Our findings evidence that SAP dynamics are less affected by orthostatic stress than by cognitive load elicited by the execution of an arithmetic task. Physiologically, this can be ascribed to the augmented involvement of upper brain centres in controlling the vascular dynamics and resistance associated with sympathetic activation. Relatively complex pattern of vascular resistance changes (an increase in splanchnic and skin regions associated with its decrease in limbs) results in an augmented SAP dynamical complexity, as demonstrated by previous works [20–22,63,66]. The trend towards lower SAP complexity values during tilt may be related to the synchronization of peripheral vascular activity due to sympathetic activation, contributing to regularize the fluctuations of SAP [67].

Comparing the entropy measures obtained through the two estimators, it is possible to infer that the values obtained by *knn* are generally lower than those obtained using the *lin* approach, especially with regard to RR and in rest and mental stress conditions. The reasons of such a difference are difficult to explain and may be related to several factors, ranging from local nonlinearities or nonstationarities, which could play a role in a univariate setting like in our case, to the effect of a bias in the nonlinear measures obtained using the *knn* estimator, due to the difficulty of working on high-dimensional spaces [68]. However, despite this bias, for the majority of measures both estimators exhibit concordant changes and are equally able to distinguish between rest and stress conditions. There are three exceptions, in which the *knn* is unable to detect differences, while *lin* does, i.e. CE for RR comparing M vs R2 (Fig. 4(c.4 vs c.2)), and DE for RR comparing M vs R2 (Fig. 4(b.4 vs b.2)) and CE for SAP comparing T vs R1 (Fig. 5(c.3 vs c.1)). This finding is difficult to explain, given that the strong nonlinear dynamics contributing to short-term HRV and cardiovascular variability [20,24,69,70] are detected by *knn* estimator but neglected by model-based parametric approach. The augmented discriminative capability of linear estimator, even if due to the non-linear dynamics present in the phenomenon but not properly taken into account, may be even in perspective used in practical application for a more accurate and fast classification between rest and stress conditions [22,41]. This is also reinforced by the very low computational times required for the *lin* estimator to compute the entropy-based measures on 300-sample series length (ST standard), which is 24 times lower if compared to *knn*.

### 4.3 Ultra-short term versus short-term analysis

The main focus of this work was to assess whether using shorter heart rate and SAP time series allows to obtain the same physiological information extracted with ST series, discussed in the previous Subsection 4.1 and 4.2. Regarding the reliability of using UST RR time series to discriminate between stress and rest conditions, our time-domain results (Fig. 2) demonstrate that overall it is possible to make use of 60-sample recordings to detect the presence of either postural or mental stress compared to a rest condition. This is true despite the fact that statistically significant differences between UST and ST series are detected in R2 with regard to MEAN and in T with regard to SDNN even for 240-sample time series (Fig. 2(a.2) and (b.1)). Therefore, our results suggest that, while UST analysis implies a significant deviation of the analyzed metrics from their ST level, such deviation does not impair significantly the capability to detect the response to stress even working with shorter time series.

The above-discussed results are reinforced by correlation analysis, which reported squared Pearson correlation coefficient always above the adopted threshold for strong correlation, with the only exception being SDNN computed for N=60. Analogous remarks can be made starting from Cohen’s *d* analysis between stress and rest conditions, with similar values for all time series lengths, with only a noticeable decrease for N=60. Such results are in agreement with previous studies in the literature on RR series reporting a good agreement between UST and ST both under physical stress [32,71] and mental stress [31,72] conditions. However, the agreement decreases during the execution of a task or in presence of a stressful event that carries dynamicity in the control mechanisms [35,49,73], similarly to the trends reported in our results with regard to squared Pearson correlation coefficient. A number of studies applying UST analysis to physical stress conditions focus on the following recovery phase, showing that the dynamics are strongly influenced by the intensity of the task and the response time of SNS and PNS [32,74]; this may explain the statistically significant differences found in R2 with regard to MEAN, being R2 a post-postural stress rest.

A quite common finding in previous works is that SDNN index exhibits a lower agreement if computed through UST RR series [49,75], and this is confirmed by our results analyzing the trend of the correlation coefficient (Fig. 2(b.1) and (b.2), bottom subplots). On the other hand, the agreement is higher with regard to RMSSD index (Fig. 2(c.1) and (c.2), bottom subplots). This finding appears to be directly related to the definition of metrics, since whereas SDNN reflects RR total power, the RMSSD is instead related only to fastest variations (i.e. vagally-mediated ones) observable even from shorter time series [17]. Although it is not possible to refer to previous studies, the results obtained with regard to SAP (Fig. 3) can be discussed similarly to RR. In particular, results highlight the capability of using UST SAP time series to discriminate between stress and rest conditions up to N=60 (with just the exception of STD for postural stress), even if statistically significant differences are reported between UST and ST distributions for MEAN during T and for most conditions for STD. Similarly to RR, a very high squared Pearson correlation coefficient is reported between UST and ST distributions, decreasing with N and going below threshold for N **≤** 120 for STD. Likewise to RR, the agreement of STD measure is lower for UST series, also confirmed by the lower effect size between rest and stress conditions.

Our results confirmed the feasibility of employing UST series to carry out computation of regularity and complexity measures (see Figs. 4 and 5), already previously reported for CE and Approximate Entropy [31,41,76]. Results evidence that, apart a couple of exceptions, the significant differences between a stressful and the preceding rest condition reported using 300-sample recordings are also retained using shorter series up to 60-sample duration for all the analyzed metrics (SE, DE, CE). This result suggests that the variation of cardiovascular dynamics and of complexity produced by a phase transition can be properly assessed even using shorter records. Similar remarks were made in [21] leading to the use of a smaller number of samples for the definition of the embedding vector able to describe the past history of a system. Even if less investigated with respect to time and frequency measurements, some studies have been interested in feasibility of HRV UST analysis using information-domain indices extracted from the ECG [31,35,77] and photoplethysmographic (PPG) [78] signals. Although these studies have focused on other non-linear measures for the analysis of the predictability (e.g., Shannon Entropy [31,35,78], dynamics (e.g., Permutation Entropy [77,78], Distribution Entropy [78]) and complexity (e.g., Approximate Entropy, Sample Entropy [31,35,77,78]), their findings may be considered in agreement with ours. In fact, most of them agree that a recording of at least 3 or 2-minutes is necessary in order to have good reliability with respect to ST analysis. Our results with regard to correlation analysis on entropy-based measures have evidenced as well that the agreement between UST and ST distributions is overall very good (above threshold) for N>=120 for RR and for N>=180 for SAP, except for non-parametric SE that exhibits a severe decrease of *r*^2^ already from 4-minutes length recordings. Therefore, we may suggest that when considering UST heart rate variability it is possible to decrease even more the time series length if compared to blood pressure variability analysis. Another interesting finding is that the correlation coefficient computed on entropy-measures during mental stress (Figs. 4 and 5, right panels) appears in most cases higher if compared to the R2 condition, similarly to what was achieved in [31] using a different validation tool.

The Cohen’s *d* analysis on entropy measures evaluated on RR evidenced a better discrimination of postural stress than mental stress with a higher effect size (Fig. 4, central subplots), in contrast to what evidenced instead with regard to SAP (Fig. 5, central subplots). In any case, the effect size decreases when shortening the time window length, thus suggesting a lower discriminative capability between stress and rest states caused by the information loss about slower dynamics due to the shorter time series.

Finally, as regards the comparison of estimation methods for entropy-based measures, the same considerations made in Subsection 4.2 hold also for the UST analysis. For the majority of measures both estimators are similarly capable of distinguishing between rest and stress conditions. There are the same three exceptions discussed with regard to ST series, in which the *knn* is unable to detect differences while *lin* is, i.e. CE for RR comparing M vs R2 (Fig. 4(c.4 vs c.2)), and DE for RR comparing M vs R2 (Fig. 4(b.4 vs b.2)) and CE for SAP comparing T vs R1 (Fig. 5(c.3 vs c.1)). This may be due again to the significant proportion of nonlinear dynamics contributing also to UST HRV and cardiovascular variability [20,24,69] detected by *knn* estimator but neglected by model-based parametric approach. Also in this case, the increased discriminative capability of linear estimator and its lower computational costs may be exploited for discrimination between rest and stress conditions [22]. Furthermore, our results demonstrate that using shorter time series also requires reduced computational costs for both estimators, with a decrease of ∼1.7 times for *lin* and ∼3.1 times for *knn* when shortening the time series length from 300 to 120 samples. Computing all the information indices exploiting time series of 120 samples through the parametric estimator is ∼11 times less computationally expensive than using the model-free estimator.

## 5. Conclusions

In this work, a comparison between ST and UST analysis has been carried out by computing physiological indices in time and information-theoretic domain on heart rate and blood pressure variability time series, during rest and both orthostatic and mental stress conditions, to assess to what extent UST analysis can represent a valid substitute of the ST gold standard, especially for stress detection.

Our results showed that time-domain and entropy-based measures computed on RR and SAP series are able to discriminate between rest and stress even for very short time series length, up to N=120 or even N=60 samples in most cases. However, the drop in correlation below the set threshold reported for UST shortest windows (N **≤** 120) suggests caution in the use of very-short time series segments, especially when analyzing SAP variability, opening a new issue that deserves further investigation in the future.

Finally, the comparison between a more reliable but time-consuming model-free estimator and a linear model-based approach suggests that the latter can be suitably employed for detecting changes in physiological conditions, thanks to the considerable benefits in terms of reduction of computational costs, at the expense of information loss about non-linearities in cardiovascular dynamics.

The combined use of UST series and faster linear estimators for entropy-based measures can be beneficial for integration of such metrics within wearable devices for a real-time monitoring of cardiovascular parameters. A future extension of this work could focus on performing the same UST analyses on photoplethysmographic (PPG) signals which can be directly acquired on wearable devices. Several studies agree on considering pulse rate variability assessed from PPG as a surrogate of heart rate variability [23,79,80] and recent studies have also investigated the use of PPG time series shorter than 300-samples as an alternative for the standard ST analysis [33,78]. Moreover, analyzing PPG signals allow to extract information also on blood pressure [81,82], and this would permit to achieve an insight into both heart rate and blood pressure variability which, as seen from our results, often yield complementary results.

## Acknowledgments

G.V. Ph.D. grant is supported by Istituto Nazionale Previdenza Sociale (INPS) Ph.D. fellowship, project title “Sviluppo di protocolli sperimentali e impiego di soluzioni tecnologiche finalizzate alla valutazione oggettiva e quantitativa dello stress lavoro-correlato”. C.B. is supported by the project “Sensoristica intelligente, infrastrutture e modelli gestionali per la sicurezza di soggetti fragili” (4FRAILTY), funded by Italian Ministry of Education, University and Research (MIUR), PON R&I grant ARS01_00345, CUP B76G18000220005. M.J. is supported by grant VEGA 1/0283/21. R.P. is partially supported by European Social Fund (ESF) - Complementary Operational Programme (POC) 2014/2020 of the Sicily Region.

